# Computational Fluid Dynamics in Highly Complex Geometries Using MPI-Parallel Lattice Boltzmann Methods: A Biomedical Engineering Application

**DOI:** 10.1101/2025.05.25.656026

**Authors:** Reza Bozorgpour

## Abstract

This study aims to establish predictive criteria for identifying cerebral aneurysms that are likely to respond favorably—either stabilizing or shrinking—following flow diverter stent (FDS) treatment. We analyzed the pre-treatment hemodynamics and geometry of four patient-specific aneurysms to determine features linked to positive outcomes. Blood flow was simulated using a massively parallel, in-house developed CFD code. Hemodynamic metrics—including vortex structure, velocity field, wall shear stress (WSS), time-averaged WSS (TAWSS), and oscillatory shear index (OSI)—were quantified in each case. Aneurysms that responded well to FDS showed lower OSI and elevated WSS and TAWSS near the neck region. In contrast, poor responders exhibited larger vortexes and persistently low WSS and TAWSS within the sac, accompanied by high OSI. Geometric factors, such as smaller neck and sac sizes and greater distance from the skull base, also appeared to favor a positive response. These results highlight a combination of geometric and hemodynamic parameters that may serve as effective predictors of FDS treatment success and support more informed decision-making during preoperative planning.

## Introduction

A cerebral aneurysm refers to an abnormal bulging of the blood vessel wall, commonly found at arterial bifurcations or curvatures. Rupture of cerebral aneurysms results in subarachnoid hemorrhage, a condition linked to high rates of mortality and morbidity [1]. Therefore, treatment planning is a vital stage to prevent an aneurysm rupture. However, having the capability to predict which treatment plan may work better for a patient-specific aneurysm will assist clinicians with more accurate and efficient treatment decision-making process.

Current treatments for cerebral aneurysms include surgical and endovascular options. Early approaches like Cooper’s artery ligation [2, 3], Cushing’s silver clips [4], Dandy’s neck ligation [5], and operating microscopes [6, 7], laid the groundwork for modern techniques. Innovations followed with Norlen and Olivecrona’s improved clips [8], Drake’s fenestrated clips [9], and bypass surgery by Crowell and Yasargil [10–12]. Dott’s wrapping technique [13, 14] and Guglielmi’s detachable coils, approved in 1995 [15, 16], revolutionized endovascular treatment. Balloon-assisted (BAC) and stent-assisted coiling (SAC) further expanded options [17–19]. Recent devices like the Woven EndoBridge (WEB), pCONus, and PulseRider stents provide better support for complex aneurysms [20–29]. Advances in materials, such as liquid embolic and covered stents, continue to enhance treatment success [30–32]. Among the less invasive options, endovascular treatment using a flow diverter stent (FDS) has gained widespread use, particularly for complex, wide-neck, and large aneurysms. The FDS is made up of numerous braided wires and is typically positioned within the parent artery at the aneurysmal neck. Its main role is to reroute blood flow along the direction of the parent artery, reducing blood entry into the aneurysmal sac, which promotes the gradual exclusion of the aneurysm from the cerebral blood circulation and encourages thrombosis within the aneurysm. The ability of the FDS to facilitate aneurysm occlusion is directly linked to the changes in blood flow that occur after the stent is deployed [33].

In last two decades, analyzing hemodynamic features, which are computed using CFD [34–38], has provided accurate prediction criteria to identify growing aneurysms, guided the selection of the most effective endovascular intervention for a specific patient-specific aneurysm [39–46], aided in assessing the effectiveness of endovascular therapies [47–52], helped with testing the performance of new endovascular devices [53, 54]. It is well established that the progression of intracranial aneurysms is closely associated with the hemodynamic conditions within the cerebral vasculature [39, 55–62]. Hemodynamic stress can result in endothelial dysfunction, triggering the release of inflammatory cytokines and exacerbating inflammation [63]. Shear stress exerted on the arterial walls is known to contribute to vascular remodeling, leading to aneurysm enlargement and eventual rupture [64]. Numerous studies indicate that wall shear stress (WSS) plays a key role in the initiation, growth, and rupture of intracranial aneurysms [65, 66]. Blood flow patterns and the distribution of WSS are key factors influencing the progression of intracranial aneurysms. Meng et al. [66] identified two groups of hemodynamic features that contribute to aneurysm growth, first group includes low WSS and elevated oscillatory shear index (OSI) magnitude while second group classified with high WSS accompanied by positive WSS gradients. Several other studies have also reported that low WSS causes degenerative changes in the aneurysm wall, triggering rupture through inflammatory responses [43, 65, 67, 68]. Furthermore, hemodynamic parameters linked to WSS that account for variations over the cardiac cycle are essential in evaluating intracranial aneurysm progression, rupture risk, and the effectiveness of endovascular treatments [62–63]. Moreover, indicators like OSI, endothelial cell activation potential (ECAP), and relative residence time (RRT) have been shown to be useful for monitoring aneurysm growth and assessing the impact of FDS on aneurysmal blood flow [33]. A recent clinical research analyzing 338 aneurysms identified lower pressure loss as a key risk factor for small intracranial aneurysm rupture [69]. However, to the best of our knowledge, the association between hemodynamics and aneurysm morphology and a patient’s response to treatment has not been yet studied. Our goal is to develop a massively parallel computational fluid dynamics (CFD) code, implementing the LBM and applying it to quantify hemodynamic parameters that can help us identify aneurysms responding positively to FDS. Using our in-house developed code the blood flow is simulated through four patient-specific aneurysms and hemodynamic features including blood flow behavior and the distribution and magnitude of velocity, WSS, TAWSS, and OSI are analyzed.

## 1. Methods

To explore the impact of hemodynamics on aneurysm treatment outcome, four patient-specific aneurysms were selected, focusing on variations in their location and size. This retrospective analysis utilized Magnetic Resonance Angiography (MRA) images in Digital Imaging and Communications in Medicine (DICOM) format, which were retrieved from patient records at two distinct time points, before and after treatment. We utilize Newtonian flow models in our simulation, though blood does not exhibit this behavior, and there remains no agreement on the optimal viscosity model for simulating blood flow in cerebral aneurysms [50, 70]. Some researchers suggest that blood viscosity can be modeled as Newtonian in cerebral arteries because the shear rates are believed to be high enough to surpass the point at which non-Newtonian properties would typically come into effect. [50, 71]. Several studies suggest that in medium and large arteries, blood viscosity remains nearly constant, supporting the assumption of Newtonian behavior in blood flow simulations [72]. Some studies claim that Newtonian viscosity results in excessive predictions of aneurysm wall shear stress, which could compromise the effectiveness of predicting the risk of rupture [50]. Based on the reviewed literature, we have chosen to use a Newtonian fluid model for our simulations. Despite the non-Newtonian nature of blood, the consensus in many studies is that the high shear rates present in cerebral arteries justify the use of a Newtonian model.

### 1.1. Geometry Preparation

In this study, we reconstructed image-based geometries for four patient-specific intracranial aneurysms, selected based on the availability of high-quality imaging data and documented clinical outcomes following graft stenting. Follow-up imaging confirmed the post-intervention results. These cases were not part of a formal clinical trial but represent individual clinical cases with sufficient data for detailed CFD analysis. The patients included a female (80 years old) with a fusiform aneurysm (Patient 1, AD), a male (76 years old) with a saccular aneurysm (Patient 2, OMW), a female (72 years old) with a saccular aneurysm (Patient 3, JT), and a female (71 years old) with a fusiform aneurysm (Patient 4, PV). MRA images were acquired at two stages—prior to and following intervention—and segmented using *Materialise Mimics* (Materialise, Leuven, Belgium). A segmentation approach was applied that preserved the arterial length on both the inlet and outlet sides of each aneurysm. The resulting anatomical models were exported as STL files, accurately representing the three-dimensional vascular topology for subsequent CFD analysis.

After generating the 3D triangulated surface, we manually trimmed the inlet and outlet vessels at visually appropriate locations where the vessel cross-sections appeared stable and circular. No artificial flow extensions were added. These truncated geometries were then used directly in the LBM simulations.

Figure 2 shows the geometry of each patient at two different points in time, before and after intervention. We have developed an in-house C++ code for both geometry processing and simulation. The process began by reading the STL file and determining the bounding box dimensions based on the minimum and maximum coordinates of the triangular facets, ensuring it encompassed the entire geometry. Uniformly spaced points were generated within the bounding box, and a collision detection method was employed to classify these points as fluid or non-fluid. If the number of triangles intersected by a ray originating from a point in a specified direction is odd, the point is classified as fluid; if even, it is classified as non-fluid. To enhance computational efficiency, we implemented a parallel processing approach using the Message Passing Interface (MPI). The bounding box and surface triangulation were distributed across multiple processors, with each processor responsible for a specific region (Figure 3).

**Figure 1:**
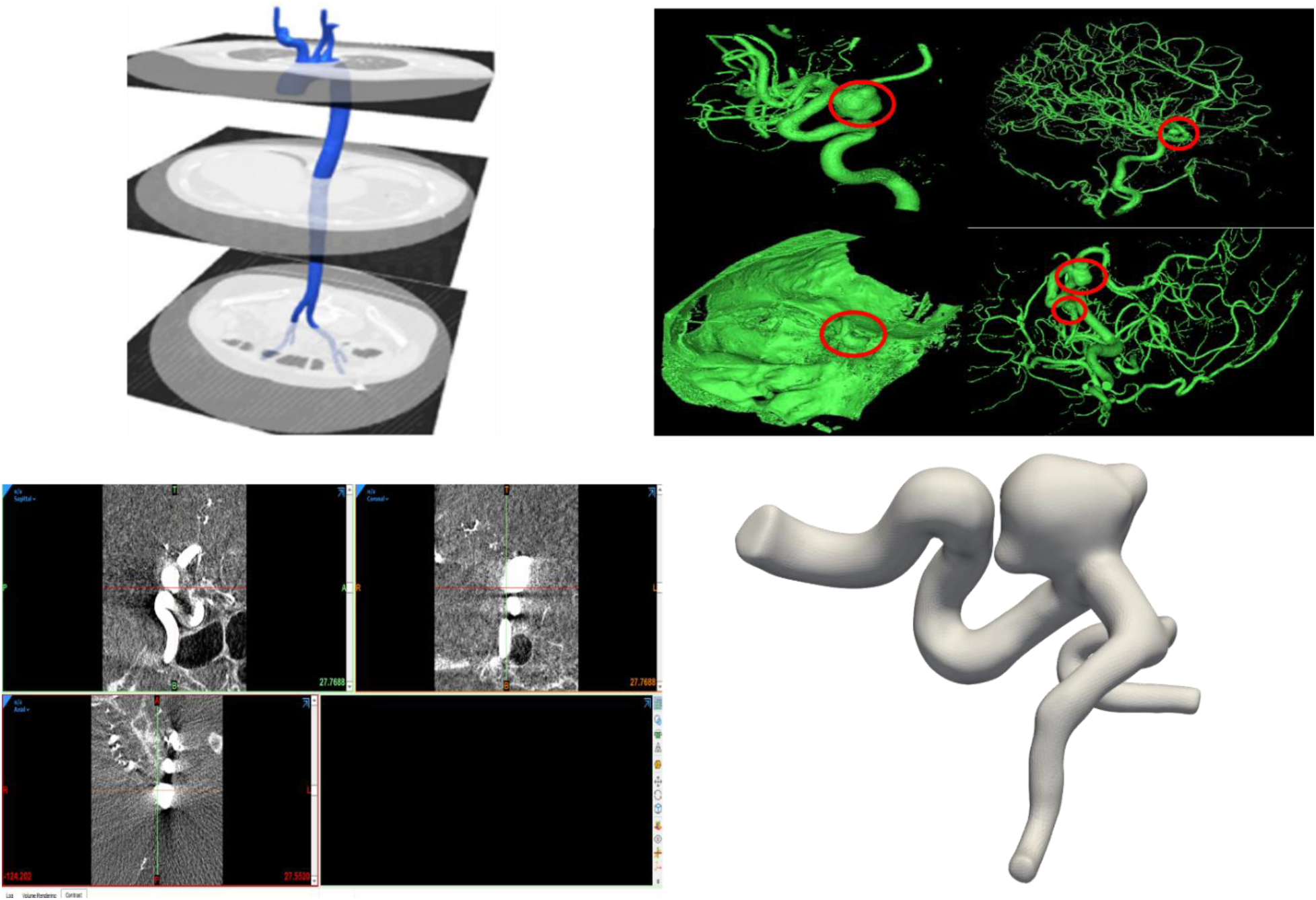
Patient-specific cerebral aneurysm reconstruction: MRA segmentation, aneurysm localization (red circles), image slicing, and final 3D geometry.

**Figure 2:**
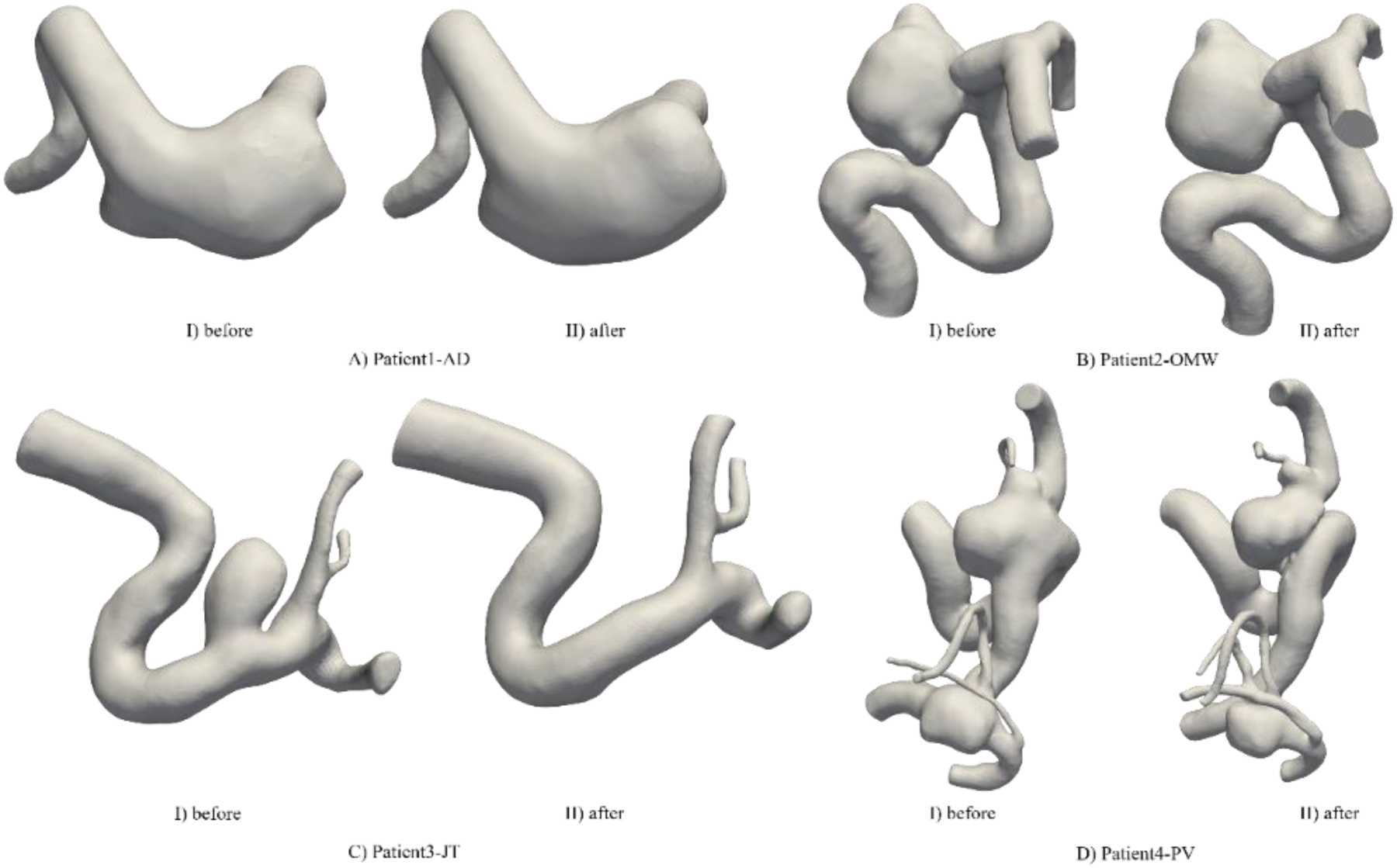
Visualization of a cerebral aneurysm before and after intervention. The left panel for each patient shows the aneurysm’s morphology prior to treatment, while the right panel presents the altered vascular

**Figure 3:**
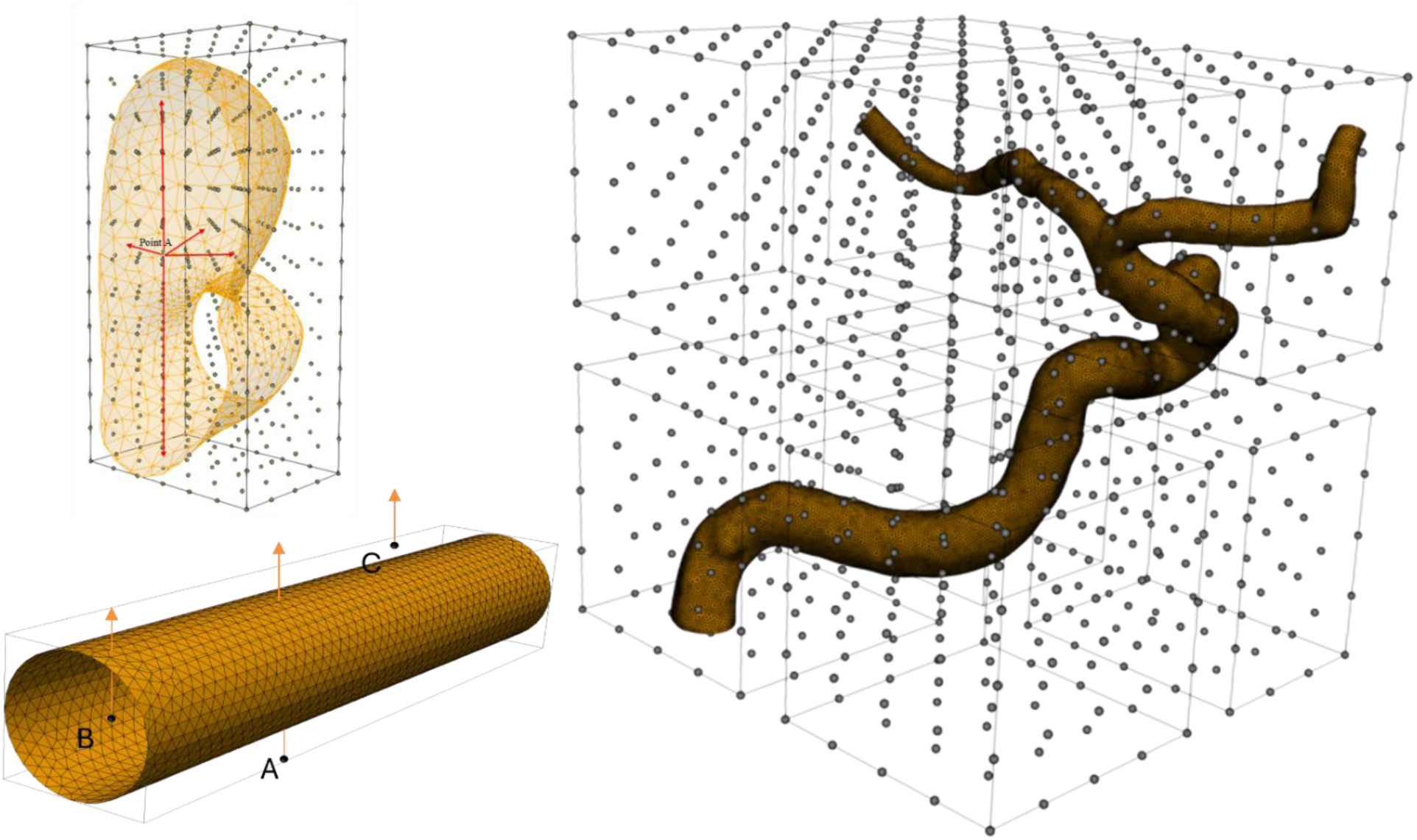
Collision detection and spatial domain decomposition for vascular geometry meshing.

A layer of ghost cells was added at the boundaries of each processor’s domain, ensuring that boundary points could be accurately identified in the presence of neighboring processors. A point was considered a boundary point if it was fluid and had at least one neighboring non-fluid point. This process was carried out on the high-performance computing cluster at the University of Wisconsin-Milwaukee, utilizing 8 AMD compute nodes. Each node was equipped with two 64-core AMD EPYC Milan 7713 processors (2.0 GHz) and 256 GB of RAM, providing a total of 128 cores per node, and 1024 cores in total. This parallel implementation significantly improved computational speed and ensured the accurate identification of boundary points within the complex geometry.

#### 1.1.1. Octree Data Structure

An Octree is a tree data structure in which each internal node can have at most eight children. Unlike a binary tree, which splits space into two segments, an Octree divides space into up to eight parts, known as octants (Figure 4).

**Figure 4:**
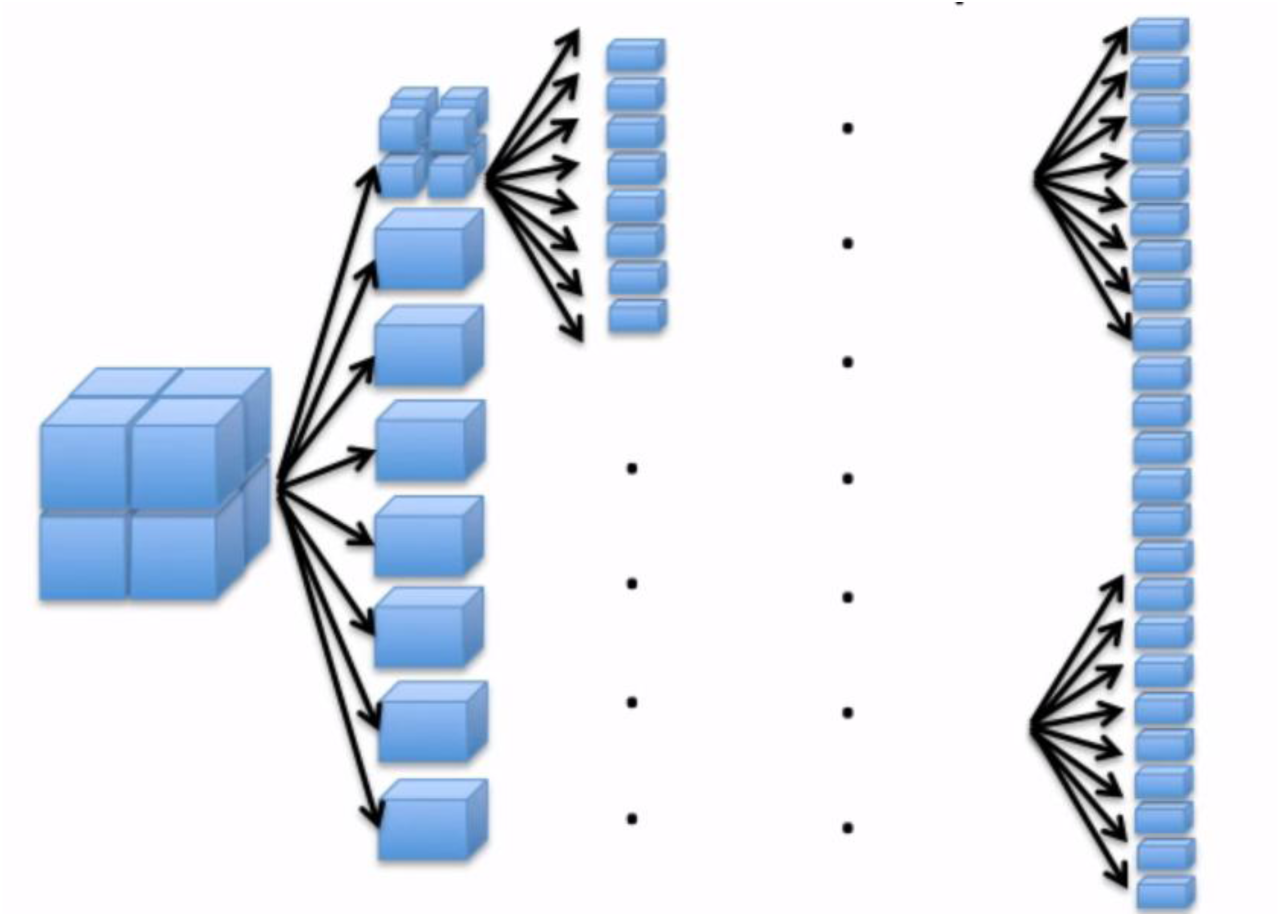
illustration of the recursive subdivision of a 3D space using an Octree data structure, where each level refines the spatial partitioning into smaller regions, optimizing data organization and computational efficiency.

It is commonly used to store 3D point data, which can require significant memory. When all internal nodes of an Octree contain exactly eight children, it is referred to as a full Octree. To construct an Octree within each MPI processor, the assigned subdomain is recursively divided into eight spatial boxes. If a box contains more than one point, it is further subdivided. The process continues until each box contains one or zero points, ensuring an optimal hierarchical representation of the data. Using this approach, instead of checking ray intersections with all surface triangles, we first restrict the computation to the specific MPI processor responsible for that spatial region. Inside each processor, the domain is divided into octants, and each octant contains only a subset of the surface triangles. Depending on the depth of the Octree, we perform ray-box intersection tests first, and then only check for intersections with triangles assigned to the intersected octants. This reduces the number of intersection tests, significantly improving efficiency. To further optimize the in-house code, points are generated one at a time within the bounding box and retained only if they lie in the fluid domain; otherwise, they are discarded. By combining MPI for parallel distribution and Octree for efficient spatial partitioning, this method enables fast and scalable ray casting, making it feasible to generate evenly spaced points inside a cerebral aneurysm while efficiently handling large datasets in a high-performance computing (HPC) environment.

#### 1.1.2. Boundary Classification

The points are classified as fluid or non-fluid in the previous section. The fluid points are further categorized into inlet, outlet, wall, and internal fluid points. To achieve this, we use the following approach: if a fluid point has a neighboring non-fluid point in any mesoscopic direction, it is labeled as a boundary point. The remaining fluid points are classified as internal fluid points. Among the boundary points, those closest to the inlet or outlet planes are labeled as inlet or outlet boundary points, respectively. The rest of the boundary points are designated as wall points. Figure 5 demonstrates this categorization of points in the 2D complex geometry.

**Figure 5:**
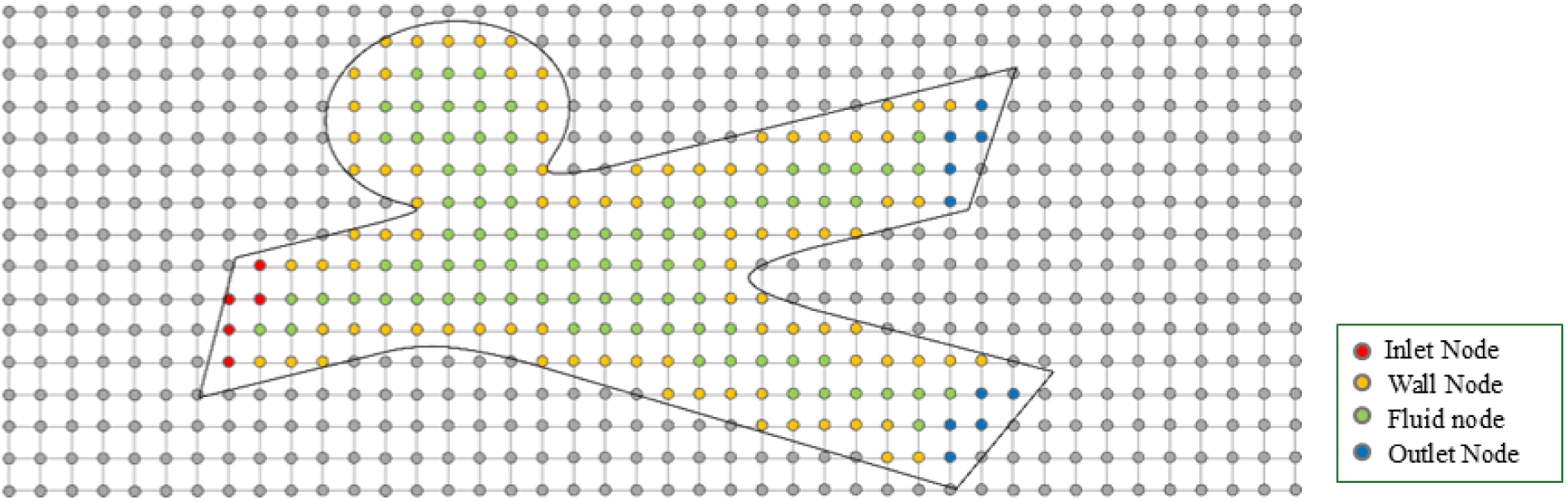
2D analogy of the node tagging process

### 1.2. Lattice Boltzmann Method

The LBM evolved from lattice gas automata and works on a discrete version of the linearized Boltzmann equation using a regular Cartesian grid (Eq. *1*). Unlike Navier-Stokes methods that target macroscopic flow properties, LBM focuses on the mesoscopic scale, dealing with the particle distribution function. The discretized Boltzmann equation is

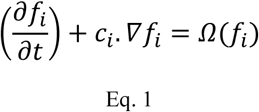

The Bhatnagar–Gross–Krook (BGK) model is widely used due to its simplicity and computational efficiency, making it well-suited for many applications, including the present study. This model simplifies the adjustment of distribution functions towards an equilibrium state 𝑓^𝑒𝑞^through a linear relaxation method, governed by the relaxation factor 𝜏. The relaxation factor 𝜏 is directly related to the kinematic viscosity 𝜈 by the formula 𝜏 = ^𝜈^ + 0.5 where 𝑐_𝑠_ denotes the speed of sound within the lattice, typically 𝑐^2^ = ^1^⁄ for standard isotropic Cartesian lattices. While the BGK model has known limitations, such as potential numerical instability as 𝜏 approaches 0.5, these issues are well-managed within the range of parameters employed in this work. The model strikes an effective balance between accuracy and computational cost, which is essential for the large-scale simulations conducted in this study. In the LBM, 𝑓_𝑖_(𝑥, 𝑡) indicates the likelihood of a particle being at position 𝑥 and time 𝑡, with velocity 𝑐_𝑖_ from a set of chosen discrete velocities. The collision term on the right side of Eq. 1 models the rate at which the fluid state transitions toward thermodynamic equilibrium. The BGK model is the most commonly used formulation for this collision term [73].

Eq. 1 describes a partial differential equation for the density function 𝑓_𝑖_. It’s usually solved by an explicit time discretization process involving two steps: collision and streaming, applied at each grid point 𝑥:

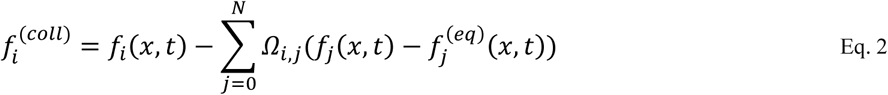

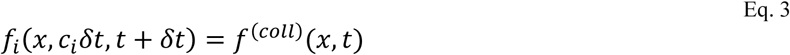

The equilibrium distribution 𝑓^𝑒𝑞^ depends solely on the fluid velocity 𝑢 and density 𝜌. The collision matrix Ω includes parameters that govern how the distributions relax towards this equilibrium state. Various formulations exist for both 𝑓^𝑒𝑞^ and Ω based on the selected collision model. For instance, in the BGK model, Ω is represented as 𝜏𝐼. The macroscopic fluid properties, such as velocity 𝑢 and pressure 𝑃, are derived from the density functions 𝑓_𝑖_.

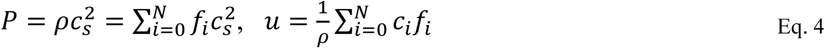

Discrete velocities 𝑐_𝑖_correspond to lattice structures, connecting each node 𝑥 to its neighbors 𝑥 + 𝑐_𝑖_. For 3D fluid simulations, lattice structures typically use 15, 19, or 27 velocities. We use the BGK collision operator for its simplicity but higher 𝜏 value to avoid instability issue with a 3D lattice structure comprising 19 velocities [74].

### 1.3. Boundary Condition

This section outlines the approach for implementing the curved wall boundary condition within the framework of LBM. To numerically solve the Lattice Boltzmann equation, it is essential to establish suitable boundary conditions for the distribution function, 𝑓_𝑖_, using the known macroscopic variables at

each boundary.

#### 1.3.1. No-slip boundary conditions

The approach for handling curved wall boundaries in the three-dimensional LBM builds upon the method proposed by Bouzidi et al. [75]. In this implementation, the nodes adjacent to the wall boundary are categorized into three types: ’fluid nodes’ within the flow domain (F), ’boundary nodes’ located near the wall (B), and ’solid nodes’ outside the flow domain (S). These categories are illustrated in Figure 6 and Figure 7, represented by black circles for fluid nodes, gray circles for boundary nodes, and squares for solid nodes. For a boundary node, N, the populations of F, B, and S are described as follows:

**Figure 6:**
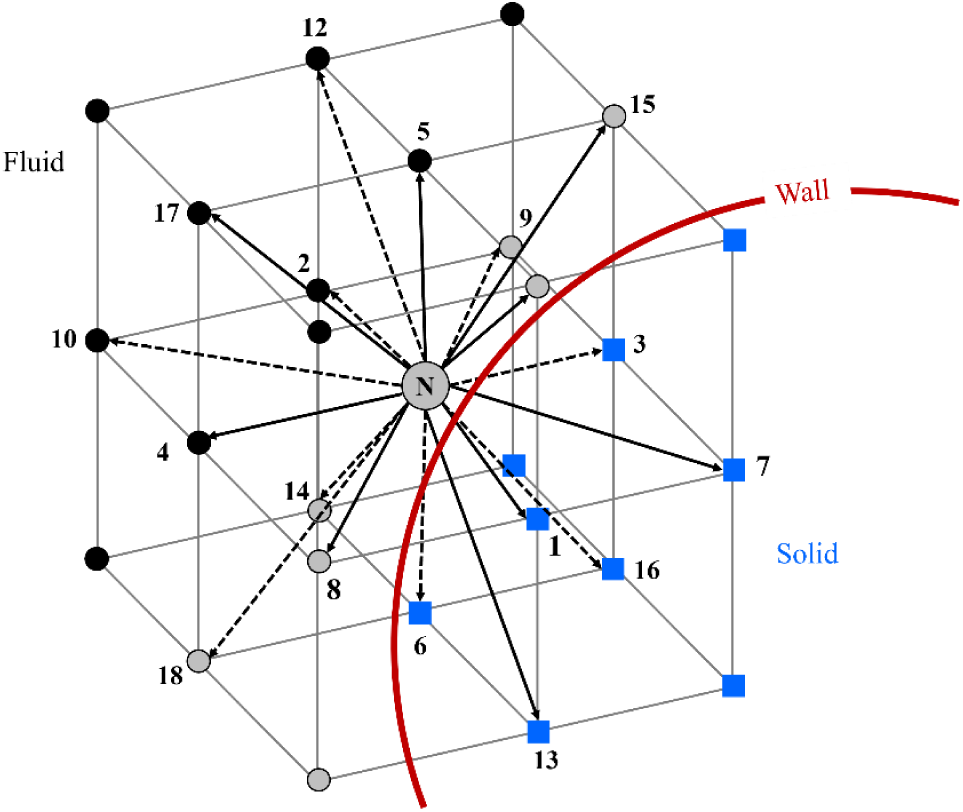
Detailed depiction of the boundary node N along with its neighboring nodes.

**Figure 7:**
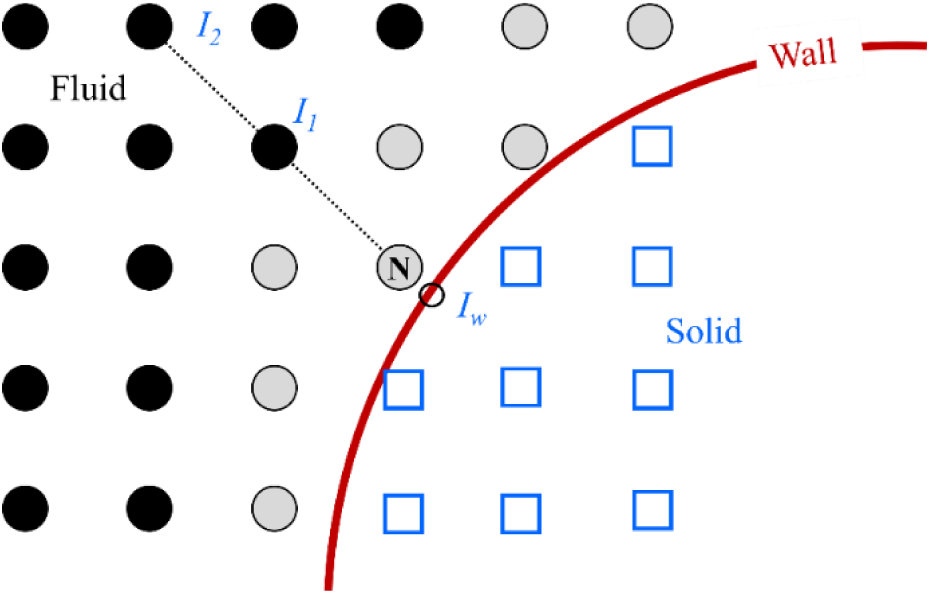
Estimation of macroscopic parameters at boundary point N using an interpolation scheme along the dashed line.

𝐹 = {2,4,5,10,12,17} 𝐵 = {0,8,9,11,14,15,18} 𝑆 = {1,3,6,7,13,16}

For nodes close to boundaries, where neighboring nodes fall outside the fluid area, unknown 𝑓_𝑖_ values are required for the streaming process. Typically, the bounce-back method is used, which assigns these unknown values by reflecting them from the opposite direction, thus simulating a particle bouncing off a wall.

However, its first-order accuracy may limit its ability to capture complex flow behavior near curved boundaries. To address this, the Bouzidi boundary condition is employed, providing higher accuracy and better handling of fluid-wall interactions in curved regions. After the streaming step, the boundary node 𝑁 has unknown populations 2,4,5,10,12,17, as these originate from nodes outside the flow domain. As a result, these correspond to the indices of the solid nodes on the opposite side, identified as 𝑆^𝑜𝑝𝑝𝑜𝑠𝑖𝑡𝑒^ = 𝑓_2_, 𝑓_4_, 𝑓_5_, 𝑓_10_, 𝑓_12_, 𝑓_17_. The macroscopic variables, such as velocity (𝑢) and density (𝜌), are known at the fluid nodes after streaming, and the wall boundary condition is applied accordingly.

To accurately implement the wall boundary condition in the simulation, it was necessary to identify which distribution functions at the wall boundary nodes needed modification. To achieve this, we constructed a vector of pairs that maps these adjustments. Each pair consists of two elements. The first element is a vector identifying the distribution functions that need modification in their opposite directions, ensuring that the distribution reflects the wall boundary condition accurately. The second element of each pair contains the fractional distance for each specified direction. This fractional distance as depicted in Figure 8 and Figure 9, is defined as the ratio of the distance from the boundary node to the intersection point with the solid wall (𝑑_𝐼𝑤_ − 𝑑_𝑁_), divided by the lattice link length (𝑑_𝐼2_ − 𝑑_𝐼1_).

**Figure 8:**
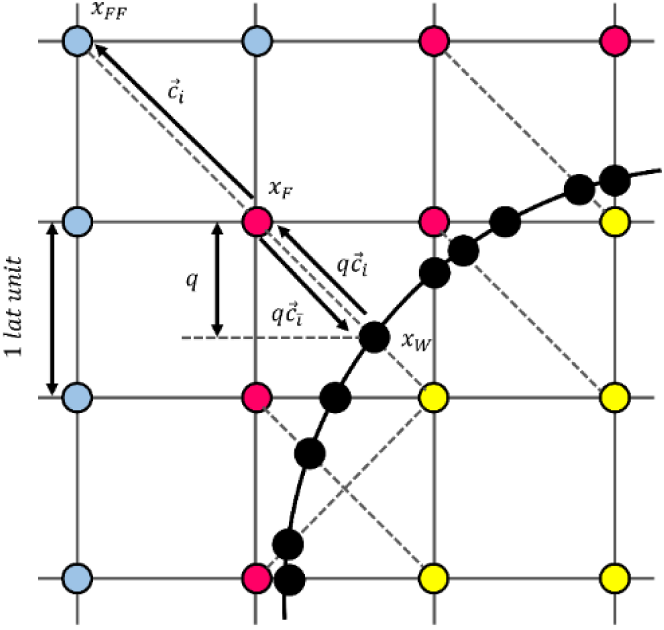
2D representation of boundary nodes, normalized distance 𝑞, discrete lattice velocities 𝑐⃗𝑖, links (dashed lines), and boundary intersections (black circle). Δ𝑥 is the spatial step.

**Figure 9:**
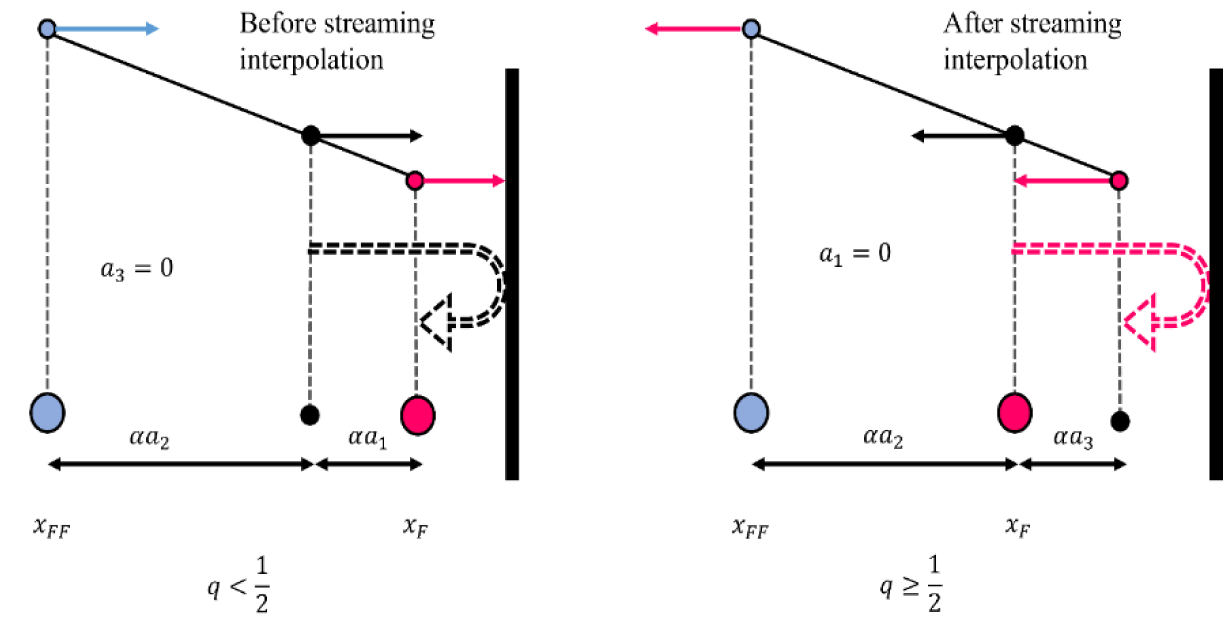
1D link-wise representation of the bouncing-back procedure in the Bouzidi method

It is crucial for interpolating the distribution functions at the boundary, as it scales the corrections based on how close the boundary node is to the wall. By implementing this mapping approach, the wall boundary condition is enforced precisely, maintaining accurate flow behavior near the wall and ensuring realistic dynamics throughout the simulation. The Bouzidi boundary condition in LBM adjusts the distribution functions at boundary nodes based on the fractional distance 𝑞 between the fluid node and the solid wall. It uses two different approaches depending on whether 𝑞 < 0.5 or 𝑞 ≥ 0.5

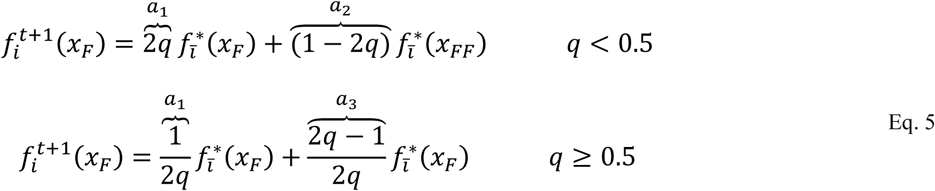

#### 1.3.2. Zou He boundary condition

In this simulation, Zou He boundary condition is used [76, 77] for both the inlet and outlet flow. For Zou He boundary condition, boundary planes can be generally described using the normal vector 𝑛, tangential vectors 𝑡_𝑖_ = 𝑐_𝑖_ − (𝑐_𝑖_. 𝑛)𝑛, and bounce-back direction 𝑓_−𝑖_ with 𝑐_−𝑖_ = −𝑐_𝑖_. From the population 𝑓_𝑖_ directed towards the wall (𝑐_𝑖_), the new populations 𝑓_−𝑖_ are calculated.

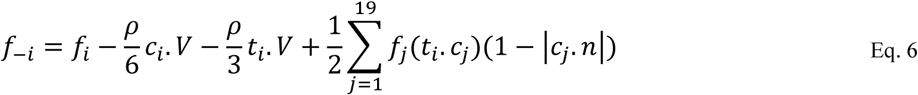

At the outflow boundary, where pressure is specified but velocity is unknown, the non-equilibrium extrapolation method is used. This method updates boundary 𝑓_𝑖_ values by extrapolating data from nearby locations.

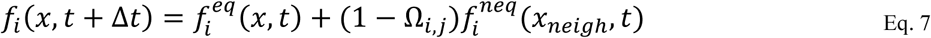

Here, 𝑓^𝑛𝑒𝑞^ = 𝑓_𝑖_ − 𝑓^𝑒𝑞^ represents the non-equilibrium portion of the distribution functions, and 𝑥_𝑛𝑒𝑖𝑔ℎ_ is 𝑖a neighboring fluid node situated along the normal to the boundary surface.

### 1.4. Pulsatile flow

Figure 10 illustrates the time evolution of the volume flow rate (VFR) measured in milliliters per minute (ml/min), relative to the reference time point H0 (in milliseconds) [89]. The gray lines represent individual raw MRI data curves, showing variability across multiple measurements. The black solid line indicates the ensemble average, providing a smoothed representation of the overall trend in VFR. The black dashed line highlights the feature point average, emphasizing key moments or averages of interest. This figure demonstrates the temporal dynamics of VFR, capturing both individual variability and averaged trends in response to the physiological event at H0.

**Figure 10:**
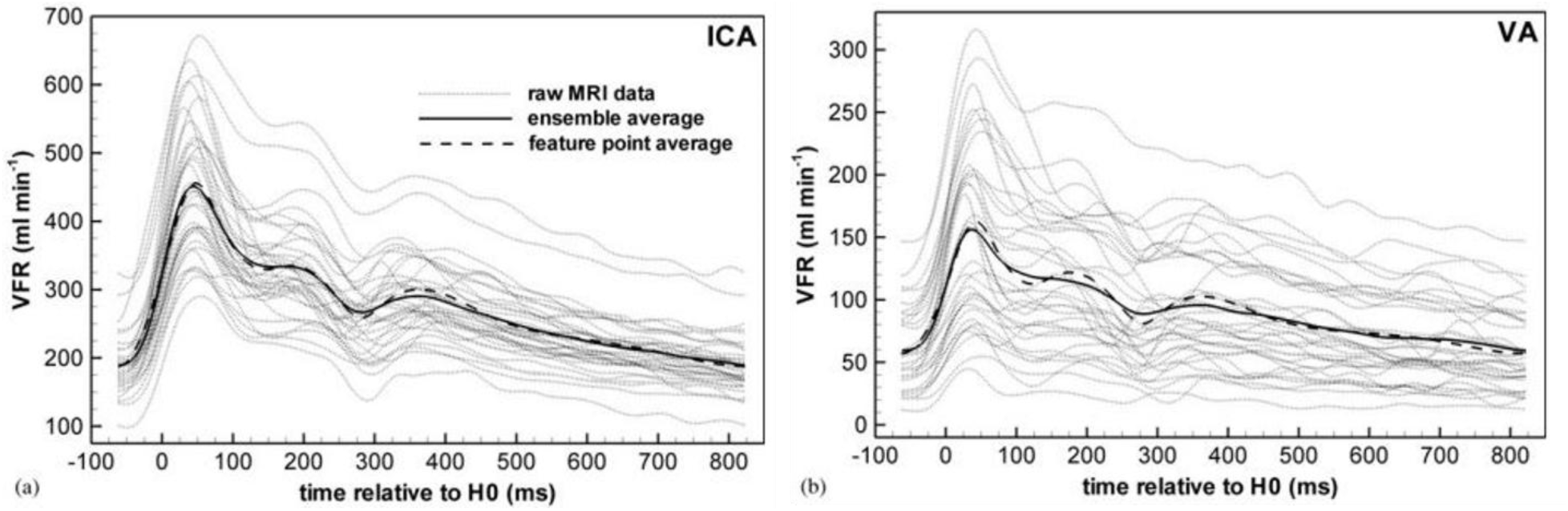
Volumetric flow rate variations over time relative to reference point H0, adopted from [78]

### 1.1. Parallelization

In an LBM simulation, steps alternate between streaming and collision. Initially, fluid particles are distributed and moved to adjacent lattice points, with values stored in a temporary function. The collide function reads temporary function, processes collisions, and updates the original array towards equilibrium. This method acts like a stencil code, updating values based on neighbor information, but uses data from a different phase space in temporary function. By using MPI, we can efficiently parallelize this process, distributing computations across multiple processors to handle larger simulations and improve performance. To update the collide function, data from the temporary array of neighboring processors is needed, which can create communication bottlenecks. To address this, ghost cells or halo cells are added (Figure 11), expanding the distribution array by one in each direction. At each step, neighboring processors exchange their border cells and receive border data for their ghost cell regions. Border data encompasses information located at the edges and faces of each subdomain. To minimize unnecessary communication, each processor is tagged to indicate whether it will participate in data exchange. This tagging system ensures efficient communication by identifying processors that need to interact. When creating fluid points within each subdomain, an important optimization is applied: if a neighboring processor in a specific direction contains no fluid points, communication in that direction is bypassed. This approach significantly reduces communication overhead, enhancing overall computational efficiency. Figure 12 represents edge and face data exchange. This process involves the transfer of distribution functions across the edges and faces of neighboring subdomains within a computational grid.

**Figure 11:**
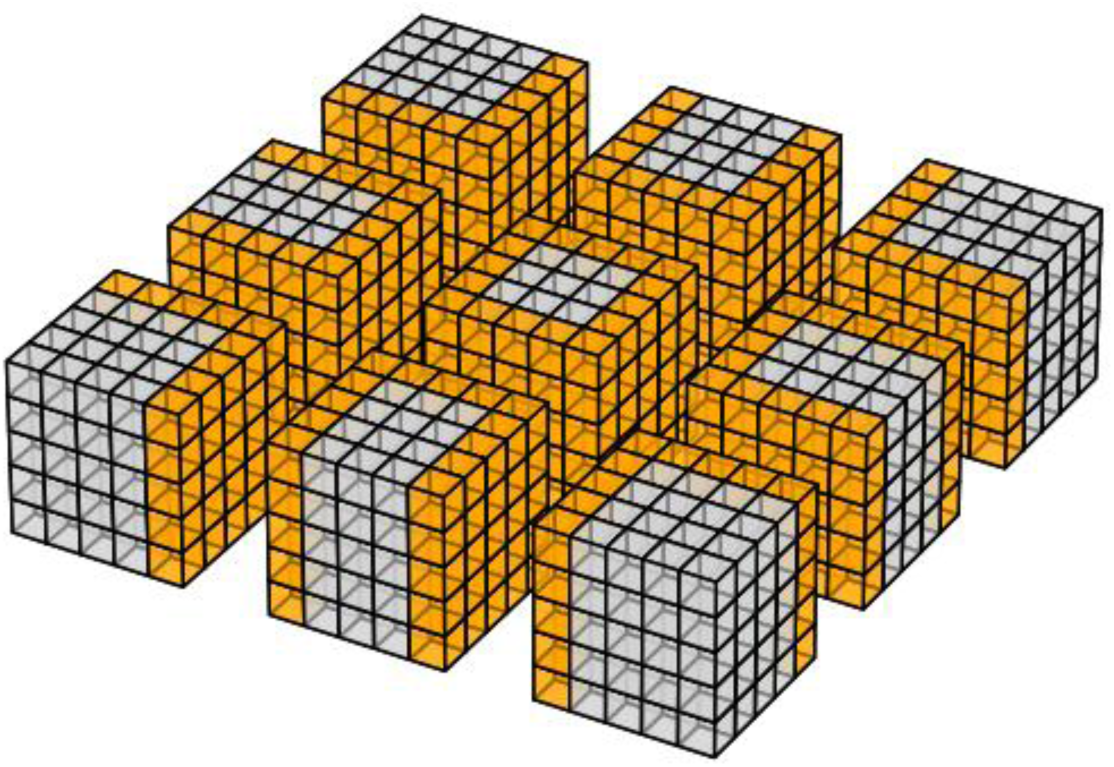
MPI 3D topology and Halo Layer representation. The figure is generated by our in-house python code

**Figure 12:**
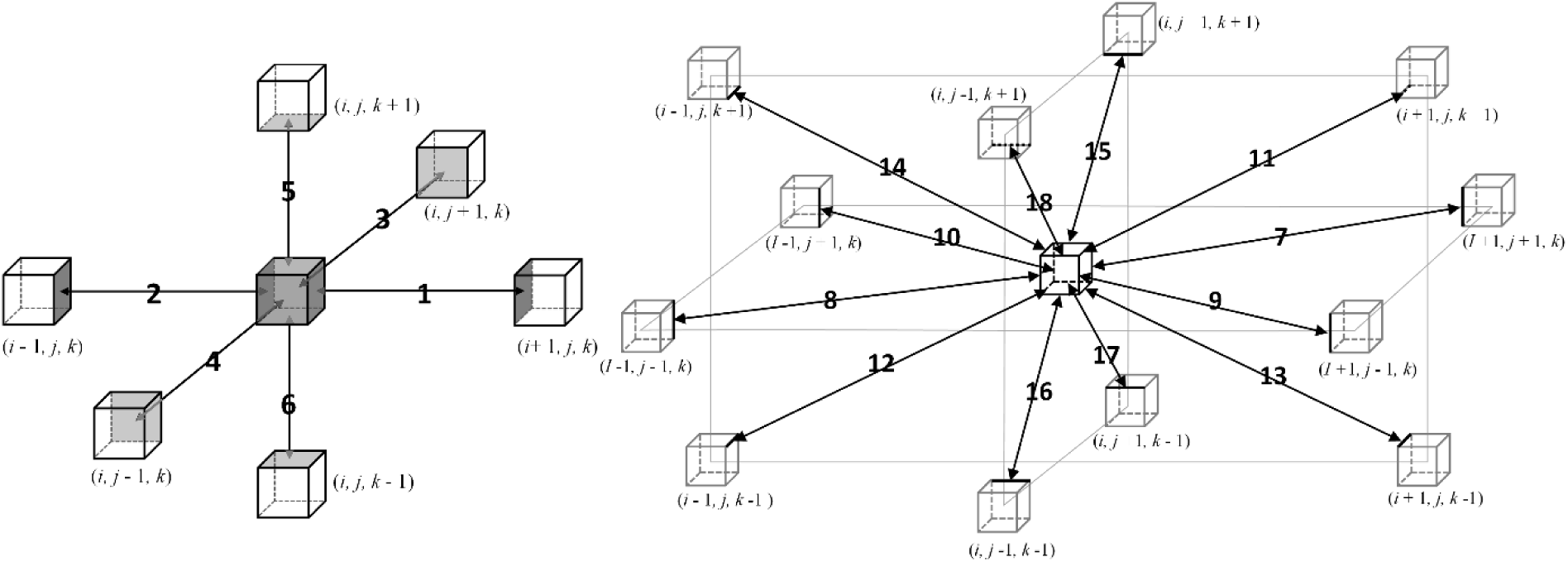
3D surface data exchange and edges data exchange

The left diagram in Figure 12, showing the 6-point stencil, represents the nearest neighbors in the x, y, and z directions, which are the nodes with which each process must communicate to share data. In the 18-point stencil on the right, the connectivity includes diagonal neighbors, requiring additional communication with more distant nodes to ensure accuracy, especially at domain boundaries. This communication ensures that all necessary data from neighboring nodes is available for the streaming step in LBM simulations. In each subdomain, we generated the mesh locally, requiring processor nodes to be designated as Fluid, Non-Fluid. The fluid node can be further subcategorized as Inlet, Outlet, Wall, or Halo. Each node represents a lattice. By knowing the inlet and outlet normal and a point on the plane, we derived the plane equation to identify

the inlet and outlet points. Wall boundary points were determined by checking if any neighboring lattices of a given lattice were non-fluid, thereby classifying it as a Wall lattice. Additional considerations were needed for lattices neighboring Halo types. The Fluid vector was constructed within each subdomain, and the collision step was applied exclusively to the Fluid lattices. However, Halo lattices were included during the streaming process.

### 1.2. Computation of Hemodynamic features

#### 1.2.1. Wall Shear Stress

When a fluid moves over a solid surface, the velocity directly at the surface is zero, adhering to the no-slip boundary condition. However, near the surface, the velocity parallel to it does not drop to zero. The change in this parallel velocity in the direction perpendicular to the surface creates wall shear stress, which is a force exerted by the fluid onto the surface. WSS can be described as the difference between the Cauchy stress on a surface and its projection in the direction normal to that surface. The stress tensor in fluids is composed of two main components: the pressure term and the viscous term.

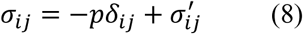

The Cauchy formula determines the total stress acting on a wall, considering the orientation of the normal vector n:

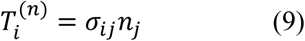

The Cauchy formula calculates the overall stress applied to a wall, factoring in the normal vector n:

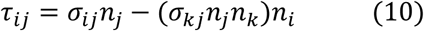

By substituting Eq (8) into Eq (10) and using the fact that 𝛿_𝑖𝑗_ 𝑛_𝑗_𝑛_𝑘_ = 1 the equation for WSS becomes:

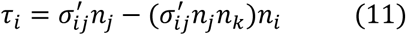

In the case of an incompressible Newtonian fluid, the viscous stress tensor σ^′^ is linearly related to the strain rate tensor 𝜀_𝑖𝑗_:

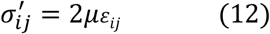

The integration of Eq (11) and (12) results in:

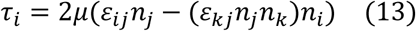

The rate of strain tensor in the LBM is defined using the non-equilibrium component of the distribution function as follows:

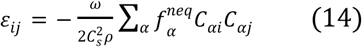

Combining equation (13) and (14) for an incompressible, Newtonian fluid, WSS in the LBM is determined using the Eq (15)

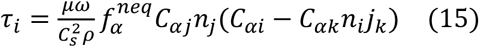

In this context, 𝐶_𝑠_ represents the lattice sound speed, μ is the dynamic viscosity, 𝜌 denotes the fluid density, and 𝜔 is the relaxation parameter in the LBM. The symbols 𝐶_𝛼𝑥_ and 𝑛_𝑥_ stand for the components of the lattice vector and the normal vector, respectively; specifically, 𝑛_𝑖_ refers to the 𝑖-𝑡ℎ component of the wall normal vector 𝑛, and 𝐶_𝛼𝑖_ indicates the 𝑖-𝑡ℎ component of the lattice vector 𝐶_𝛼_. 𝑓^𝑒𝑞^ is the equilibrium Maxwell-Boltzmann distribution and 𝑓^𝑛𝑒𝑞^ = 𝑓_𝛼_ − 𝑓^𝑒𝑞^.

#### 1.2.2. Time- Averaged Wall Shear Stress and Oscillatory Shear Index

To assess the impact of FDS placement in a patient-specific aneurysm TAWSS and OSI were computed. These time-averaged metrics effectively capture the transient behavior of blood flow and are critical for characterizing the modified hemodynamics within intracranial aneurysms post-FDS implantation. Furthermore, they provide valuable insights for evaluating device performance.

TAWSS provides the mean magnitude of the WSS vector throughout the cardiac cycle. This measure captures the overall shear stress exerted on the vessel walls over time. Here, T represents the duration of the cardiac cycle, while 𝜏_𝑤_ denotes the instantaneous wall shear stress vector at any given moment.

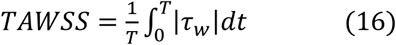

OSI is a dimensionless parameter that captures the directional variations of the WSS vector throughout the cardiac cycle relative to the primary flow direction.

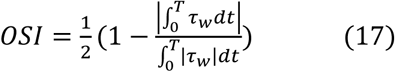

OSI values fall within the range of 0 to 0.5. A value of 0 indicates unidirectional shear stress, which is typically associated with healthy vascular conditions. In contrast, higher OSI values have been linked to triggering inflammatory responses in the artery wall.

### 1.3. Mesh independence and validation studies

To validate our code, we reconstructed a patient-specific aneurysm geometry and simulated blood flow through it using a constant inlet velocity. A mesh independence study was conducted with mesh sizes of 100, 50, 40, 30, and 20 𝜇𝑚. Blood flow velocity at a cross-section within the aneurysm stabilized at a mesh size of 20 𝜇𝑚, confirming mesh independence. Using this mesh, we compared the WSS on the aneurysm surface between our LBM code and COMSOL 6.2. The WSS distributions closely matched (Figure 13), further validating the accuracy of our LBM code in capturing complex surface hemodynamics.

**Figure 13:**
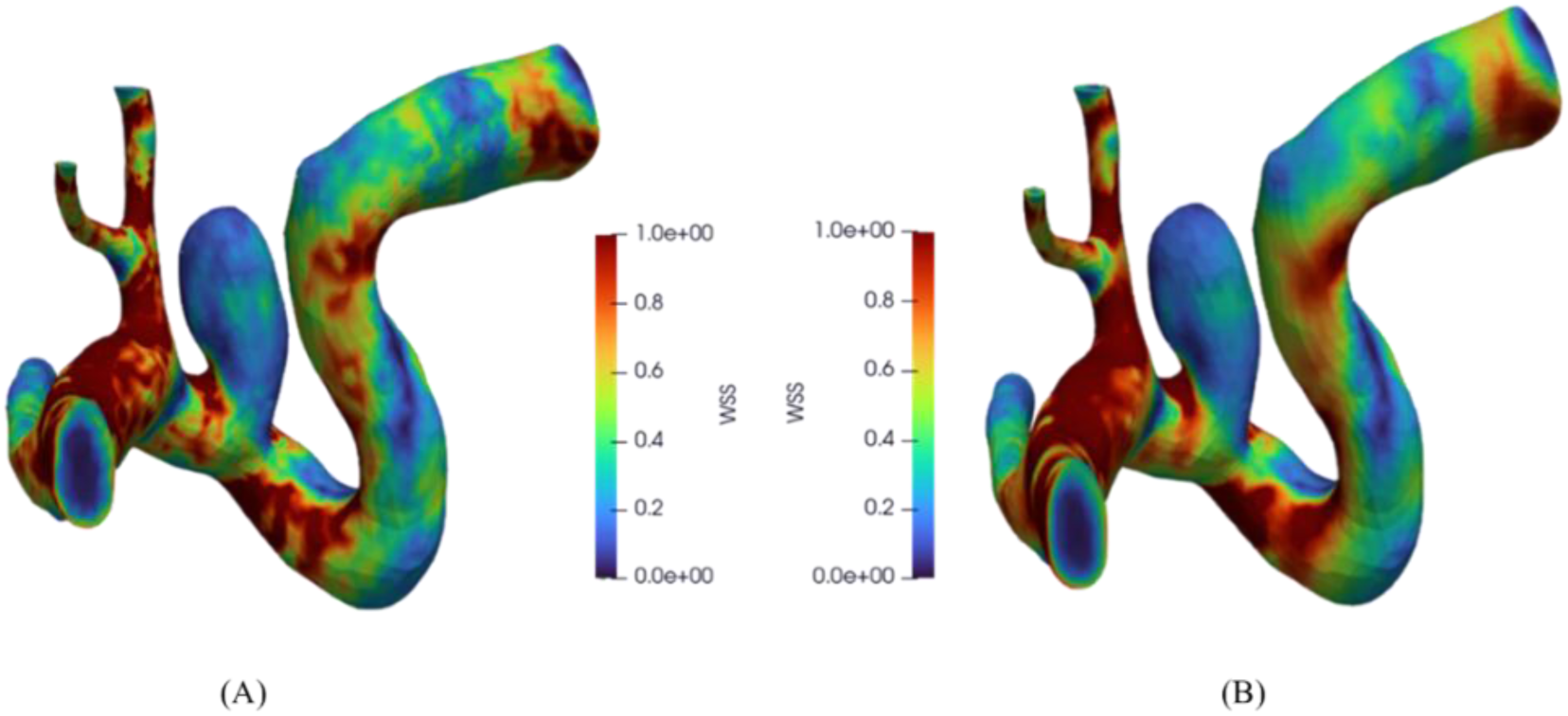
WSS distribution and magnitude for a patient-specific aneurysm. A) using our in-house developed LBM code with grid spacing 20 𝜇𝑚, B) using COMSOL Multiphysics.

The differences observed across the five grid spacings (100, 50, 40, 30, and 20 𝜇𝑚) arise from the ability of finer grids to more accurately capture flow features, reduce numerical diffusion, and improve boundary condition implementation. Coarser grids tend to underrepresent geometric details and flow characteristics, leading to deviations. As the mesh is refined, these discrepancies diminish, highlighting the importance of mesh refinement for achieving a reliable and accurate solution.

## 2. Results

This section presents a comprehensive hemodynamic analysis of four cerebral aneurysms, highlighting the velocity magnitude and wall shear stress during different cardiac cycle phases.

Figure 14 introduces the hemodynamic characteristics of Patient1-AD, including velocity and WSS at systole and diastole, along with TAWSS and OSI over the first cardiac cycle. For Patient2-OMW, Figure 15 follows the same format, presenting velocity fields and WSS patterns, as well as TAWSS and OSI distributions. Figure 16 continues this analysis for Patient3-JT. Patient4-PV, who has two aneurysms, is depicted in Figure 17 and Figure 18. Each figure illustrates the velocity, WSS, TAWSS, and OSI distributions for one of the two aneurysms. Figure 19 includes a comparative bar chart summarizing the mean values of WSS (measured at the systole of the 4th cardiac cycle), TAWSS, and OSI (averaged over the first cycle) across all four patients. These averages were calculated over the aneurysm sac, excluding regions near the parent arteries.

**Figure 14:**
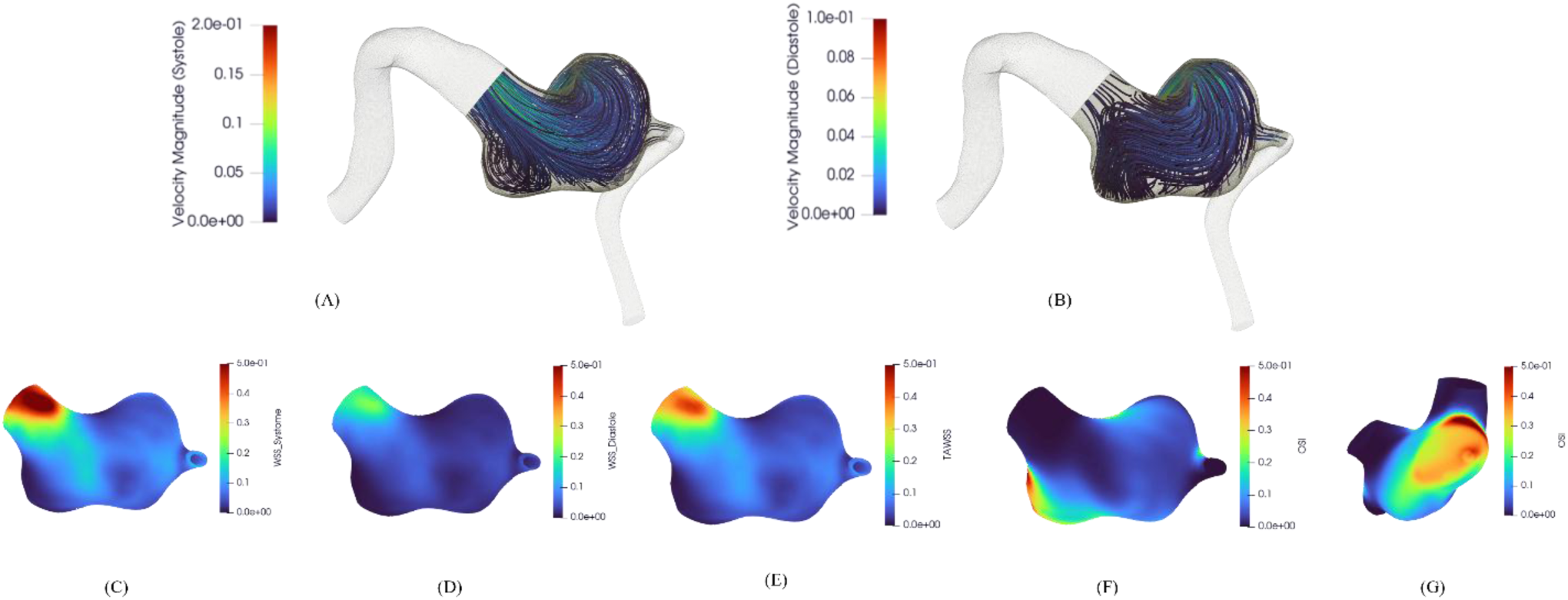
Hemodynamic assessment of Patient1-AD aneurysm. (A–B) show velocity at systole and diastole; (C–D) show WSS; (E) shows TAWSS; (F–G) show OSI from two views.

**Figure 15:**
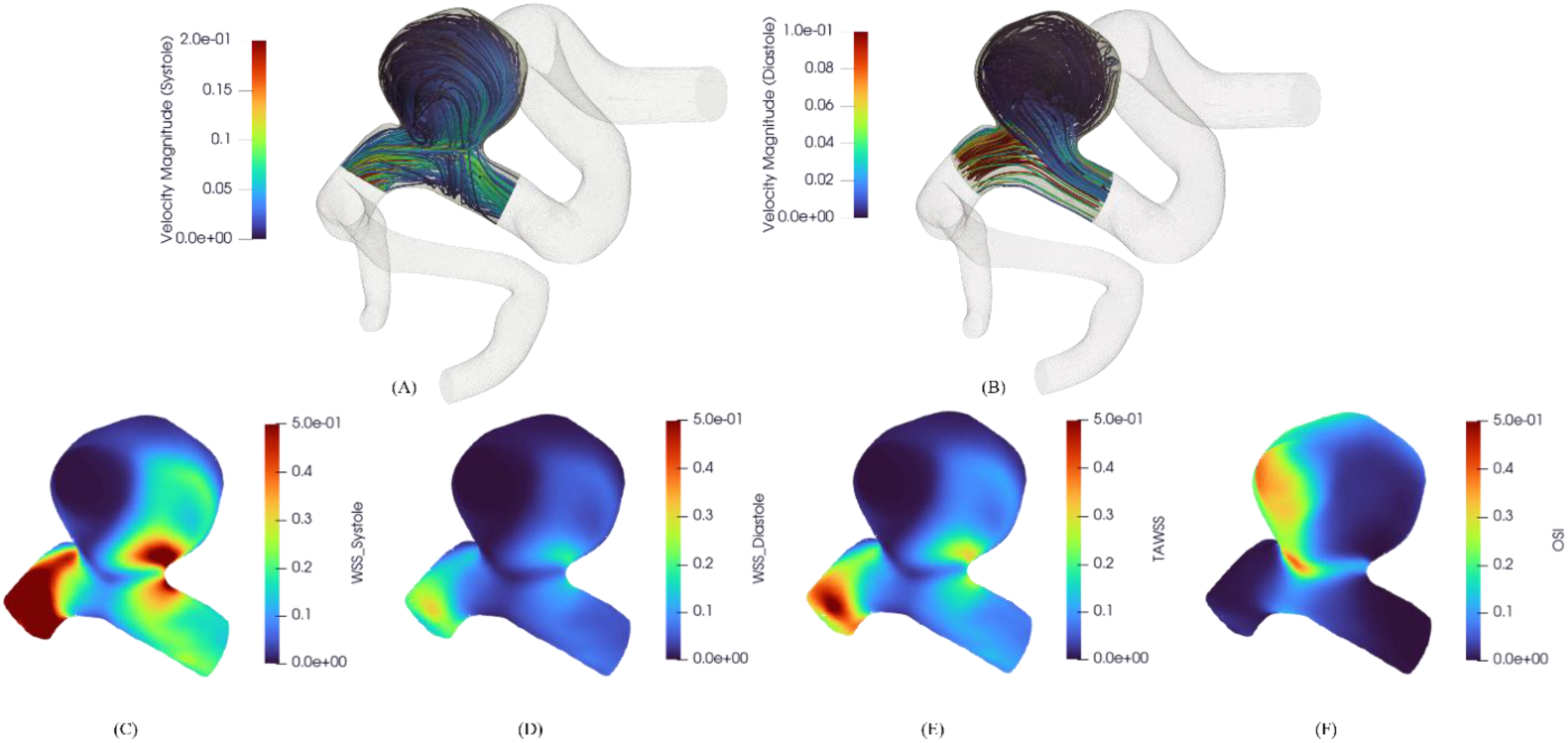
Hemodynamic analysis of Patient2-OMW aneurysm. (A–B) show velocity at systole and diastole; (C–D) show WSS; (E) shows TAWSS; and (F) shows OSI.

**Figure 16:**
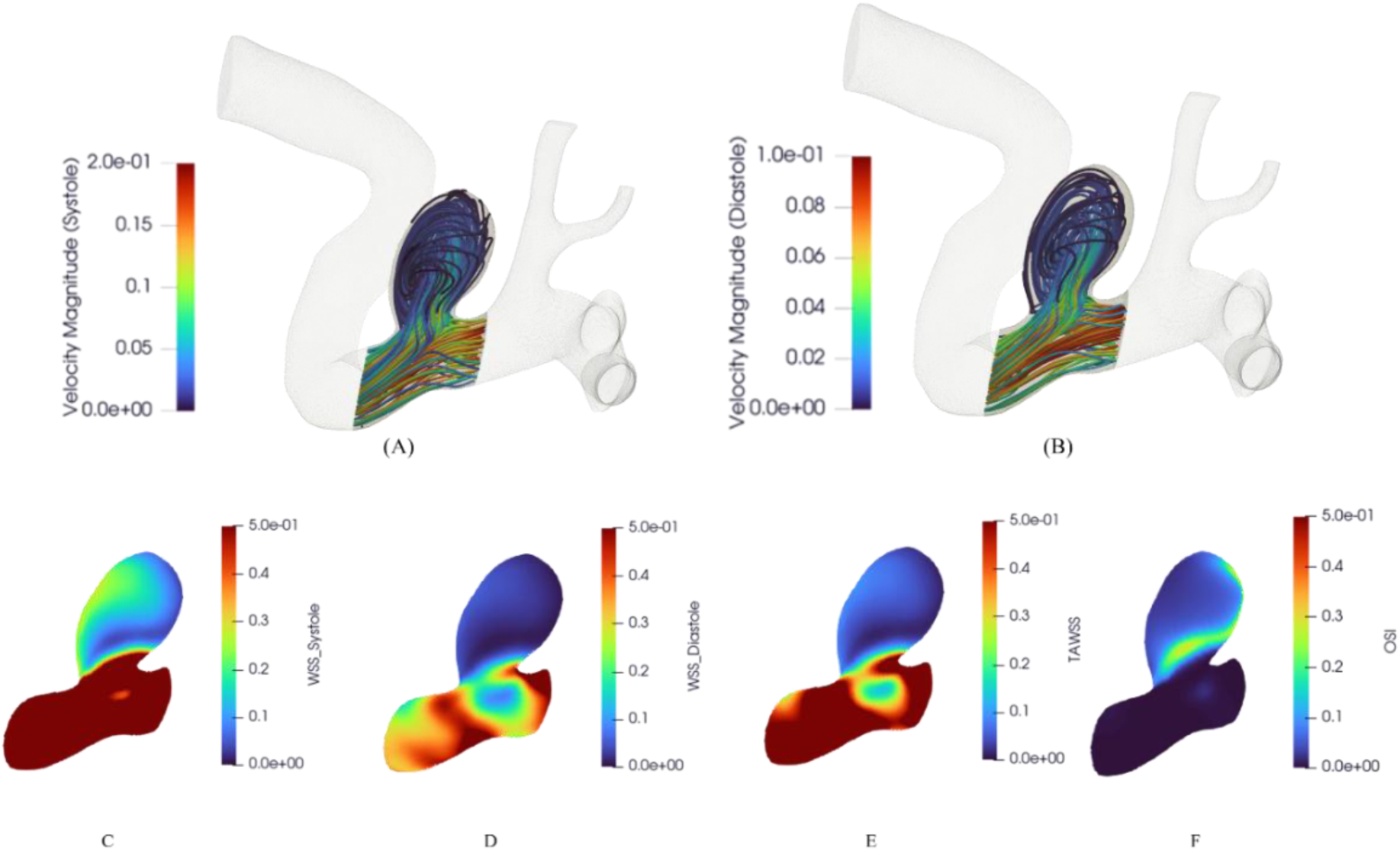
Hemodynamic analysis of Patient3-JT aneurysm. (A–B) show velocity at systole and diastole; (C–D) show WSS; (E) shows TAWSS; and (F) shows OSI.

**Figure 17:**
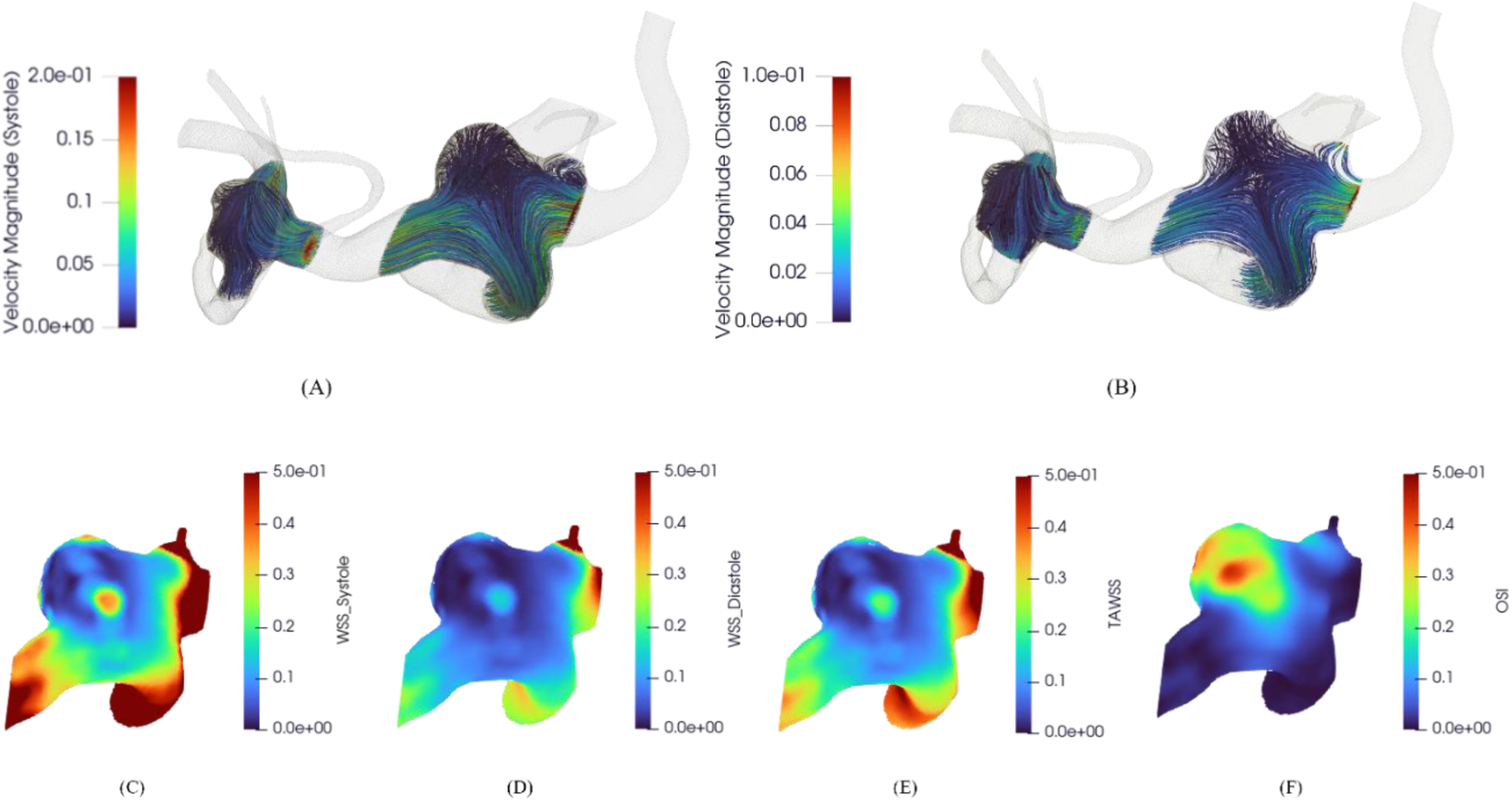
Hemodynamic analysis of a Patient4-PV with first aneurysms. A and B display the velocity magnitude and distribution at systole and diastole, respectively. C, D, E, F show magnitude and distribution of WSS at systole and diastole

**Figure 18:**
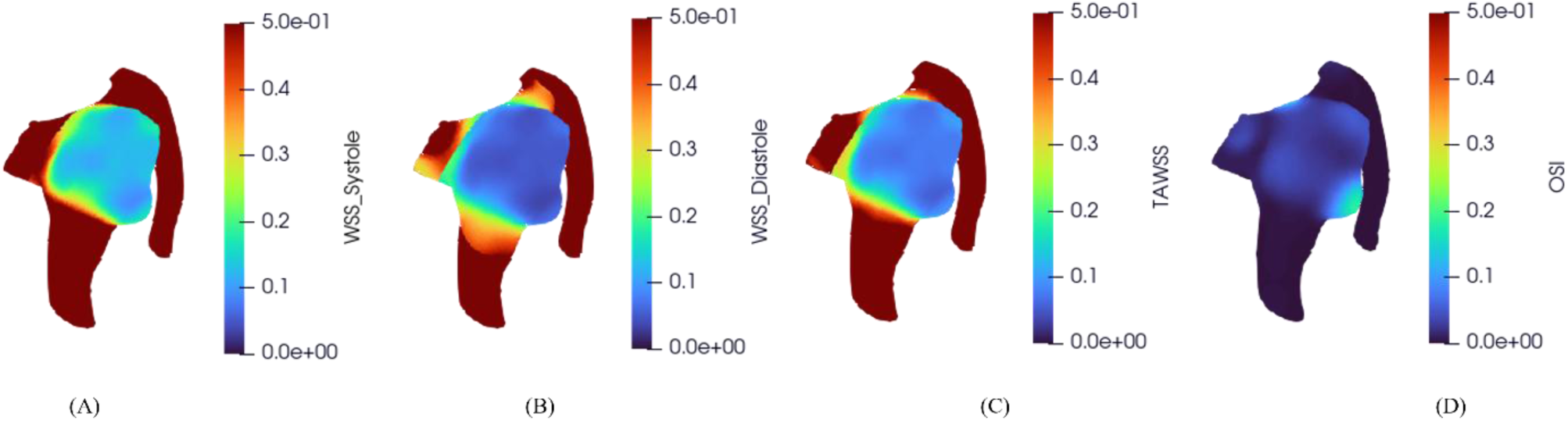
Hemodynamic analysis of a Patient4-PV second aneurysm. C, D, E, F show magnitude and distribution of WSS at systole and diastole

**Figure 19:**
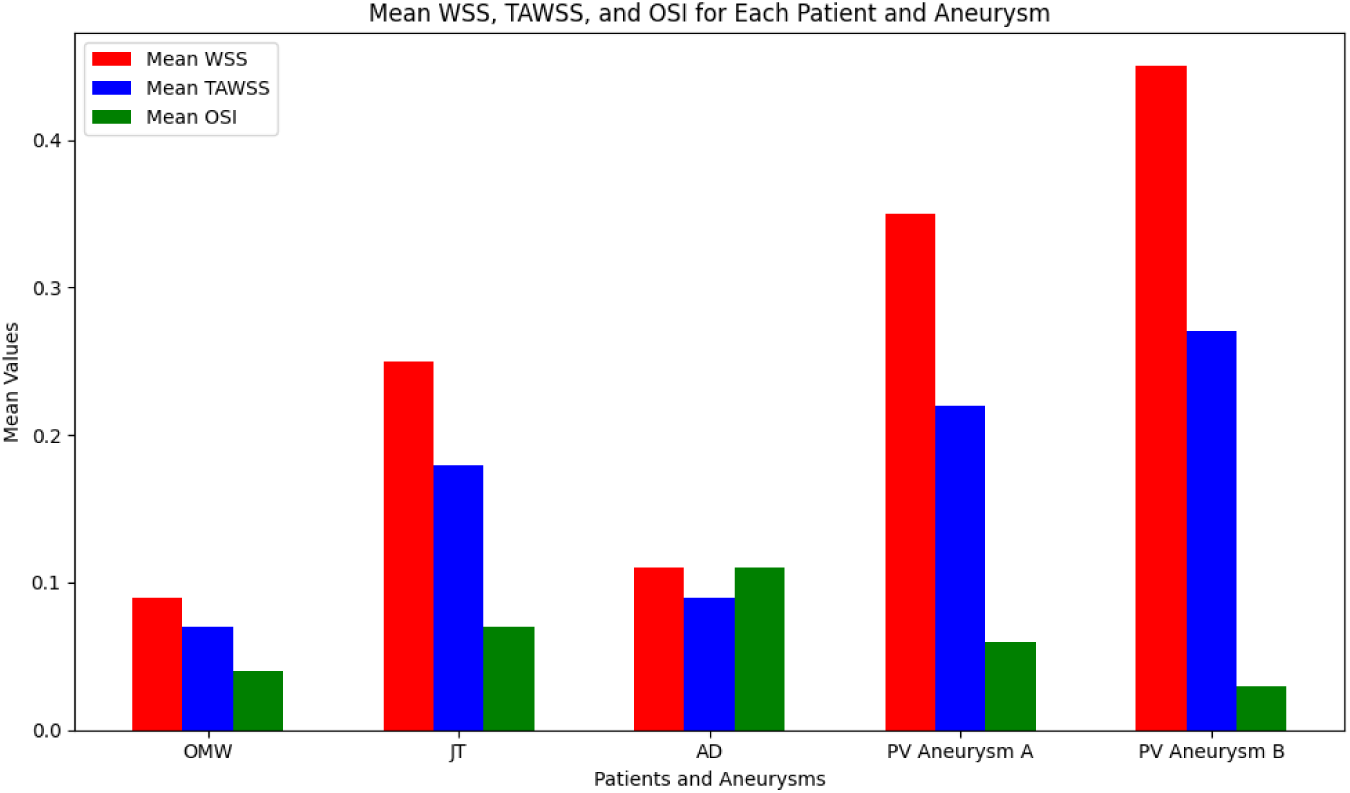
Mean values of WSS, TAWSS, and OSI averaged over each aneurysm sac.

Finally, Figure 20 provides histograms showing the distribution of WSS, TAWSS, and OSI values for each case. Panels A through D correspond to Patient1-AD, Patient2-OMW, Patient3-JT, and Patient4-PV, respectively.

**Figure 20:**
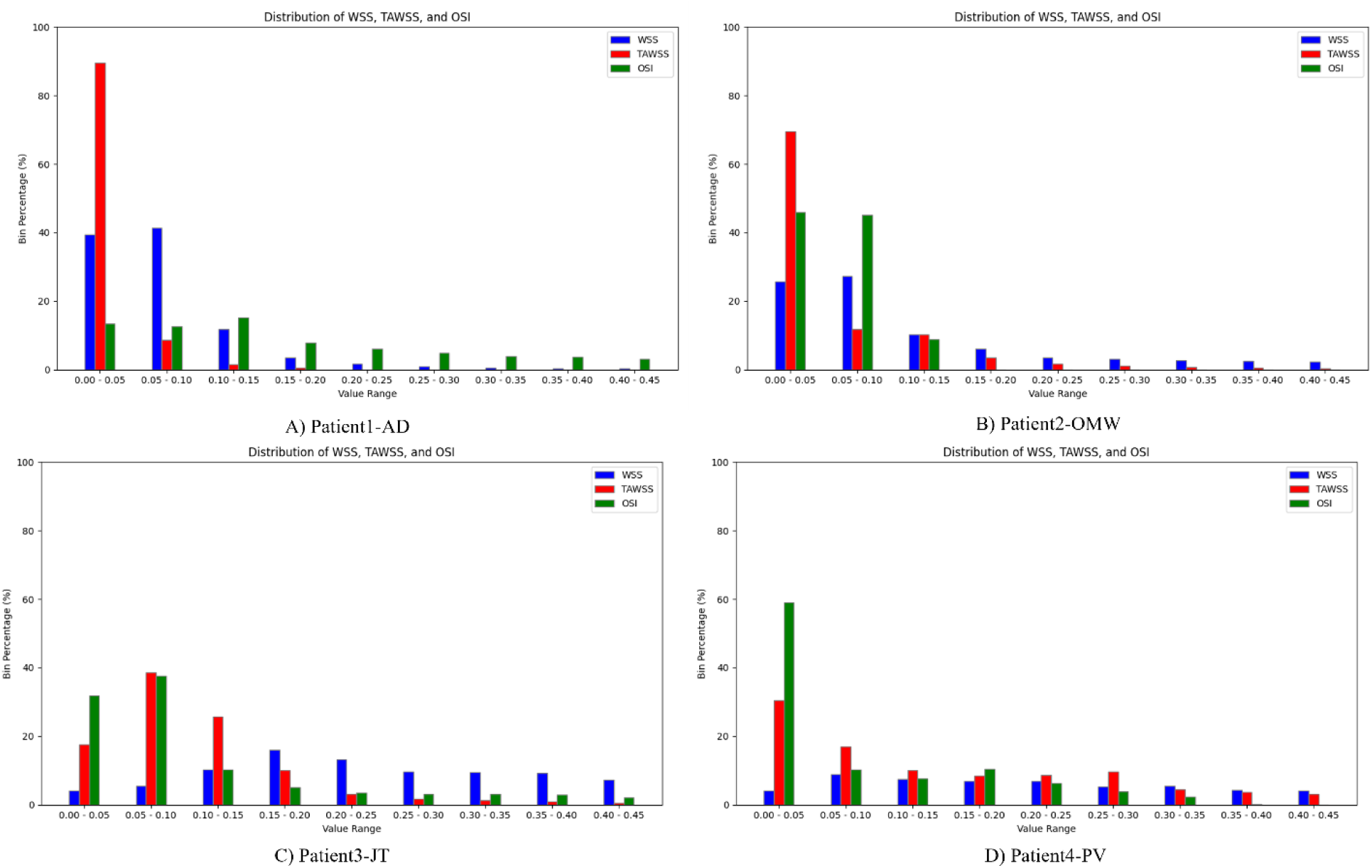
Percentage distribution of WSS (4th cycle systole), TAWSS, and OSI (1st cycle) across aneurysms of patients AD (A), OMW (B), JT (C), and PV (D).

**Table 1 summarizes the geometrical and demographic characteristics of each aneurysm case.** For each patient, we recorded the aneurysm shape, sex, and age, and measured the neck size (N) and maximum width (W) based on 3D reconstructions. These morphological parameters were used to assess geometric variability across cases and to inform the hemodynamic simulations.

**Table 1:**
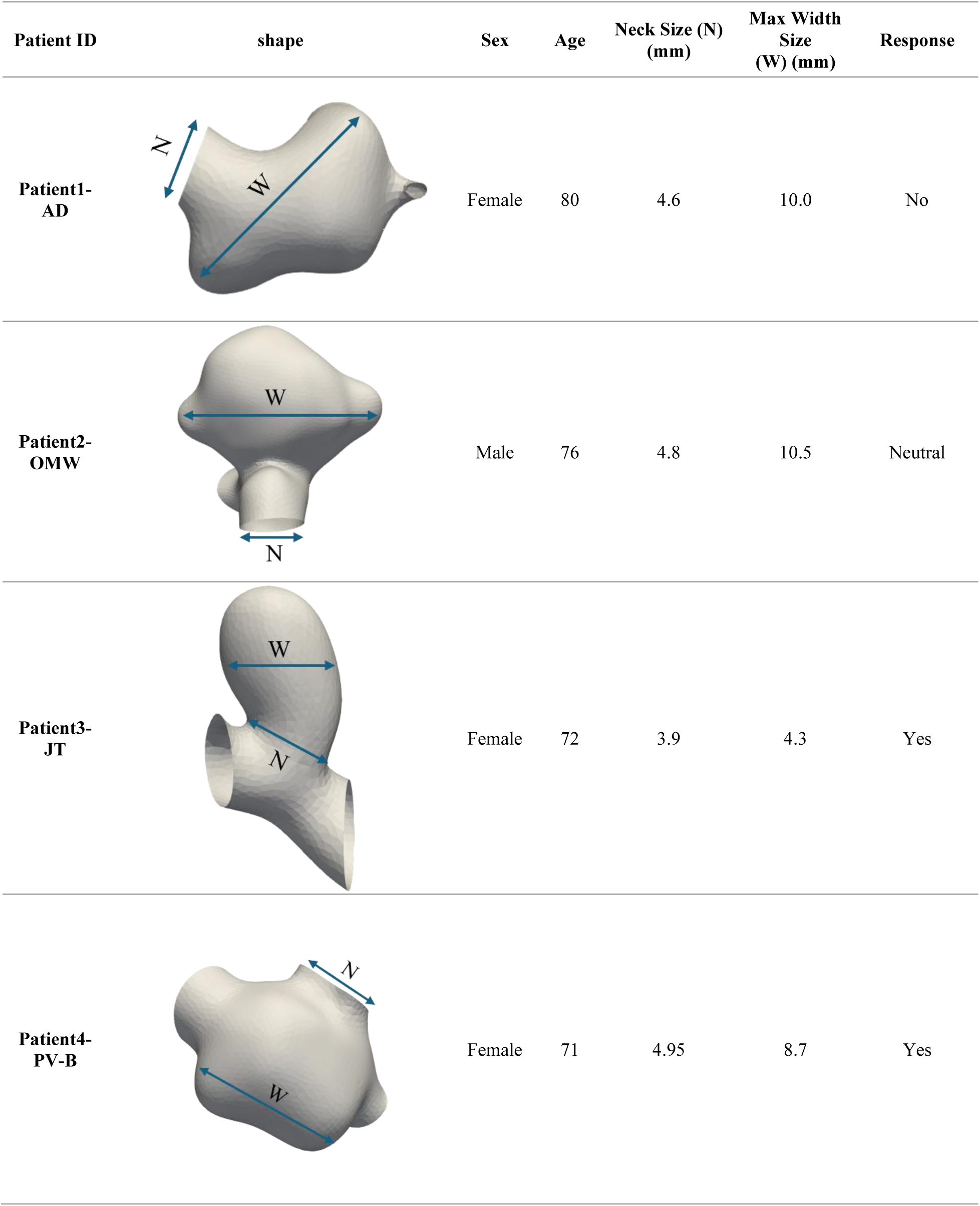
Geometrical characteristics of aneurysms.

| Patient ID | shape | Sex | Age | Neck Size (N)<br>(mm) | Max Width<br>Size<br>(W) (mm) | Response |
| --- | --- | --- | --- | --- | --- | --- |
| Patient1-AD |  | Female | 80 | 4.6 | 10.0 | No |
| Patient2-OMW |  | Male | 76 | 4.8 | 10.5 | Neutral |
| Patient3-JT |  | Female | 72 | 3.9 | 4.3 | Yes |
| Patient4-PV-B |  | Female | 71 | 4.95 | 8.7 | Yes |

## 3. Discussion

In this study, we have developed an in-house massively parallel code implementing lattice Boltzmann method to simulate the blood flow through aneurysm models reconstructed using patient-specific images. The comparative analysis of the hemodynamic factors—WSS, TTAWSS, OSI, and velocity—across all four cases sheds light on the varying responses to treatment, emphasizing why some patients experienced favorable outcomes while Patient1-AD did not. The hemodynamic investigation provides relevant information about possible influencing factors on the treatment responses. Quite striking among them is the high OSI identified in Patient1-AD, higher than in the rest of the patients, as depicted in Figure 14G. OSI refers to the amount of oscillatory, disturbed flow in a vessel and may exert a significant stress on endothelial cells and impede proper vascular remodeling. The higher value of OSI in Patient1-AD suggests that this patient probably was subjected to more disturbed flow conditions inside the aneurysm, which could result in less favorable treatment response. Oscillatory flow can adversely affect the healing process of the aneurysm wall and is associated with an increased risk for inflammation to take place. Thus, the efficacy of the intervention is also lowered. In obvious contrast, the OSI values were significantly lower in Patient2-OMW, Patient3-JT, and Patient4-PV—those who responded to the treatment favorably. This, together with the reduced OSI, would indicate more stable flow patterns, thus offering a more favorable environment for vascular healing and recovery.

With further analysis, two vortexes are observed inside aneurysm of Patient1-AD (panels A-B in Figure 14), where the blood flow appears scattered, and a few small-sized eddies or regions of disturbed flow can be viewed. This behavior will establish greater potential for adverse effects such as blood stagnation or low wall shear stress. Patient4-PV (as seen in panel A-B of Figure 17 and Figure 18) has the minimum vortex formation and represents a streamlined flow with minimal variation in the shape, which can be considered as the sign of a healthy response with minimal recirculation zones. Panels A-B of Figure 15 and Figure 16 show that in aneurysm of Patient2-OMW and Patient3-JT the vortices are at the center of the aneurysmal sac and dominant. The recirculation zone of such central vortices is well-defined, organized around a single core. The fact that this kind of vortex has a centralized structure provides stable and predictable flow patterns inside the aneurysm, contributing to the positive hemodynamic response notwithstanding significant formation of vortices.

Further differences were observed in the distribution and magnitude of TAWSS in Figures 14E, 15E, 16E, 17E, and 18C. Indeed, Patient1-AD had significant areas of much lower TAWSS compared to OMW, JT, and PV. Lower TAWSS often indicates sites where blood is either stagnant or plugs occur, which may contribute to negative vascular remodeling and impede full recovery. The relatively lower TAWSS in the aneurysm of Patient1-AD may have resulted in inferior flow dynamics, thus promoting a negative response. In contrast, the TAWSS was higher in regions of Patient2-OMW, Patient3-JT, and Patient4-PV, which means that blood flow would be more continuous and steadier. This would, in turn, support vessel stability, allowing proper function of the endothelial cells, which is very important for positive healing post-treatment. Higher TAWSS in these cases may have created a more favorable environment for successful intervention.

Previous studies have reported that high magnitudes of WSS can be considered responsible for the onset and development of an aneurysm, extended areas of aneurysm with very low WSS are associated with the potential for further formation and rupture of an aneurysm [79–83]. Patient1-AD’s aneurysm had patches of low WSS (Figure 14C) that could have contributed to continued flow disturbances and an insufficient vascular response because of the unsatisfactory treatment outcome. In contrast, Patients2-OMW, Patient3- JT, and Patient4-PV demonstrated more balanced WSS profiles, avoiding the detrimental extremes that could hinder healing [65, 66, 83–92]. Their aneurysms presented moderate WSS levels that promote vascular adaptation and repair without contributing to harmful flow conditions. Our further analysis of these cases would also suggest that, while age and gender may each have some bearing on the outcome, a balanced hemodynamic profile may be more important. For example, Patient2-OMW was an elderly male, 76 years old; Patient3-JT, a female, was 72 years old; while Patient4-PV, a female also aged 71, responded well to FDS (Table 1). This trend would thus seem to indicate that favorable hemodynamic conditions, including moderate WSS and TAWSS with low OSI, are able to allow for positive outcomes to occur, even with older individuals. On the contrary, Patient1-AD was an 80-year-old female presenting an aneurysm that continuously developed despite the assistance of FDS. Low WSS and TAWSS, together with high OSI, appeared in her profile, thus suggesting that age, combined with less stable flow conditions, may limit the efficacy of the treatment. These cases would further suggest that while age and gender may affect vascular resilience, maintaining balanced hemodynamics might be more important to ensure good treatment outcomes.

Our results for mean WSS (seen in Figure 19) reveal that Patient1-AD experienced lower WSS than Patient3-JT and Patient4-PV but not from Patient2-OMW. Despite Patient2-OMW’s positive response, the mean WSS was lower than that of Patient1-AD (0.08 Pa vs. 0.1 Pa). This indicates that WSS alone may not be the sole determinant of treatment success. The lower OSI in Patient2-OMW (0.059) compared to Patient1-AD (0.1126) may have played a more crucial role by fostering a stable flow pattern, while the higher OSI in Patient1-AD may have counteracted any potential benefits of relatively higher WSS, ultimately leading to an unsuccessful outcome. This observation underscores the complexity of aneurysm hemodynamics, where multiple interacting factors, rather than a single parameter, influence treatment success.

Histograms of WSS, TAWSS, and OSI (shown in Figure 19) in aneurysms demonstrate that Patient1-AD has a significant WSS and TAWSS in the lowest range (0.00 - 0.05), indicating that a large proportion of the aneurysmal wall experiences very low shear forces. Such low shear stress conditions are often associated with poor vascular health, potentially leading to adverse remodeling or aneurysm progression [81, 82, 93]. In contrast to the other patients, the OSI values for Patient1-AD are more evenly distributed across the entire range of values, rather than being heavily concentrated in the lowest range. This even distribution of OSI values across all intervals results in a higher overall OSI for Patient1-AD, reflecting a greater prevalence of oscillatory shear stress across the vessel wall. This higher OSI suggests increased flow disturbances, which can create an unstable hemodynamic environment hindering vascular healing. The combination of low WSS, formation of large vortexes, low TAWSS, and evenly distributed, elevated OSI might explain the lack of positive response to FDS in Patient1-AD, as these hemodynamic conditions could promote further deterioration of the vascular wall. In contrast, Patients2-OMW (panel B), Patient3- JT (panel C), and Patient4-PV (panel D) display more favorable distributions, for WSS, OSI, and TAWSS. These values extend into slightly higher ranges and OSI values are predominantly concentrated in the lower intervals (0.00 - 0.10). This distribution suggests that Patient 3-JT’s vessel wall experiences moderate shear forces with minimal oscillations, supporting a stable flow environment. Such conditions are more likely to contribute to positive vascular remodeling [94–96], which aligns with Patient3-JT’s favorable treatment response. Patient4-PV shows a broader distribution of WSS and TAWSS values that are less concentrated in the lowest range. The OSI for Patient4-PV is largely confined to lower values, indicating reduced oscillatory behavior. This low OSI concentration, coupled with moderate WSS and TAWSS distributions, points to a stable and unidirectional flow profile, likely supporting the vessel wall’s health and stability and explaining Patient 4-PV’s positive treatment response. Patient2-OMW’s profile also reflects a beneficial hemodynamic environment. Although TAWSS remains low, the WSS distribution is more balanced with a presence in higher ranges, and OSI is concentrated in lower bins, minimizing oscillations in shear stress direction. This lower OSI and moderate WSS distribution suggest a less turbulent flow, providing a stable shear environment that would support favorable treatment outcomes.

## 4. Conclusion

In this study, our main objective is to establish predictive criteria for response of a patient-specific aneurysm to FDS. This capability will assist clinicians with more efficient and precision treatment planning. We analyzed the morphology of aneurysms (location, neck width, and longest width), simulated blood flow through vasculature, and quantified several hemodynamic features determining the best predictive criteria for patient-specific aneurysm’s response to FDS. In patients with positive responses (Patient2-OMW, Patient3-JT and Patient4-PV), smaller aneurysm size, smaller neck size, being far from skull, stable flow patterns, moderate WSS near the aneurysm neck, and low OSI were observed. Conversely, the negative response in Patient1-AD was linked to significant hemodynamic disturbances, with low WSS and high OSI. This patient had the largest aneurysm among four patients, and the aneurysm was very close to the skull. These findings underscore the importance of aneurysm morphology, achieving stable flow, adequate shear near critical areas, and minimizing oscillatory patterns for successful stenting outcomes. Even though we investigated a limited number of patients, our study takes a significant first step in introducing aneurysm morphology and hemodynamic criteria to assist clinicians in identifying patients who are most likely to benefit from FDS.

## Declarations Ethical Statement

None

## Availability of data and materials

The datasets used and/or analyzed during the current study are available from the corresponding author on reasonable request.

## Acknowledgment

The authors wish to express their deep appreciation to Daniel C. Siercks, Interim Director of the UWM-High Performance Center, for ensuring seamless access to essential computational resources upon request. His support was crucial to the success of this work. The authors also extend their sincere gratitude to Pilhwan Kim and Setayesh Abiazi throughout this project.

